# TMC1 Confers a Leak Conductance to Modulate Excitability of Auditory Hair Cells in Mammals

**DOI:** 10.1101/617472

**Authors:** Shuang Liu, Shufeng Wang, Linzhi Zou, Jie Li, Chenmeng Song, Jiaofeng Chen, Qun Hu, Lian Liu, Wei Xiong

**Affiliations:** School of Life Sciences, Tsinghua University, Beijing 100084, China; IDG/McGovern Institute for Brain Research at Tsinghua University, Tsinghua University, Beijing 100084, China

**Keywords:** Cochlea, Hair cell, Mechanotransduction, TMC1, Leak, Channel, Conductance, Tonotopy

## Abstract

Hearing sensation relies on the mechano-electrical transducer (MET) channel of cochlear hair cells, in which Transmembrane channel-like 1 (TMC1) and TMC2 have been proposed to be the pore-forming subunits. Meanwhile it has been reported that TMCs regulate other biological processes in a variety of lower organisms ranging from sensations to motor functions. However, it is still an open question whether TMCs play roles other than their function in MET in mammals. In this study, we report that in mouse hair cells TMC1, but not TMC2, provides a background leak conductance, with properties distinct from those of the MET channels. By cysteine substitution, 4 amino acids of TMC1 are characterized critical for the leak conductance. The leak conductance is essential for action potential firing and tonotopic along the cochlear coil. Taken together, our results suggest that TMC1 confers a background leak conductance that modulates membrane excitability in cochlear hair cells.

## INTRODUCTION

Hair cells are mechanoreceptors that convert mechanical stimuli provided by sound and acceleration into electrical signals. In the snail-shaped mammalian cochlea, hair cells are organized into three rows of outer hair cells (OHCs) and one row of inner hair cells (IHCs) that run along the length of the cochlear duct. The cochlea is tonotopically organized, where hair cells at the base of the cochlea represent high-frequency sounds and hair cells at the apex represent low-frequency sounds with a gradient in between. OHCs amplify input sound signals while IHCs transmit sound information to the CNS.

The mechanotransduction complex in cochlear hair cells consists of a multitude of proteins including ion-channel subunits, cell adhesion proteins, myosin motors, and scaffolding proteins that are critical to sense sound-induced force (Xiong and Xu, 2018). The transmembrane proteins TMC1 (transmembrane channel-like 1), TMC2, LHFPL5 (lipoma HMGIC fusion partner-like 5 / also known as tetraspan membrane protein of hair cell stereocilia, TMHS), and TMIE (transmembrane inner ear expressed protein), are thought to be integral components of the MET channels in hair cells. TMC1 and TMC2 have been proposed to be the pore-forming subunits of the MET channel in hair cells (Ballesteros et al., 2018; Corey and Holt, 2016; Kawashima et al., 2015; Pan et al., 2018). Consistent with this model, MET currents are absent in hair cells from mice lacking both TMC1 and TMC2 (Kawashima et al., 2011), while the unitary conductance, permeability, and ion selectivity of the MET channel differs between hair cells expressing only TMC1 or TMC2 (Beurg et al., 2015a; Beurg et al., 2014; Corns et al., 2017; Corns et al., 2014, 2016; Kim and Fettiplace, 2013; Pan et al., 2013). Finally, cysteine mutagenesis experiments are consistent with the model that it is a pore-forming subunit of the hair-cell MET channel (Pan et al., 2018). However, all efforts have so far failed to express TMC proteins in heterologous cells to reconstitute ion channel function (Corey and Holt, 2016; Wu and Muller, 2016). Intriguingly, MET responses in OHCs vary tonotopically, and a lack of TMC1 and LHFPL5 but not TMC2 abolishes the tonotopic gradient in the MET response (Beurg et al., 2014; Beurg et al., 2015b). While changes in the levels of expression of TMC1 from the base to the apex have been proposed to underlie the tonotopic gradient in the MET response, the mechanisms that cause the tonotopic gradient are not completely defined (Beurg et al., 2018; Beurg et al., 2006; Ricci et al., 2003; Waguespack et al., 2007).

TMC orthologues in other species have been linked to a diversity of functions. In *Drosophila melanogaster*, TMC is expressed in the class I and class II dendritic arborization neurons and bipolar dendrite neurons that are critical for larval locomotion (Guo et al., 2016), and TMC is enriched in md-L neurons that sense food texture (Zhang et al., 2016), and for proprioceptor-mediated direction selectivity (He et al., 2019). In *Caenorhabditis elegans*, TMC-1 regulates development and sexual behavior (Zhang et al., 2015), and is required for the alkaline sensitivity of ASH nociceptive neurons (Wang et al., 2016). While efforts have failed to demonstrate that TMCs in flies and worms are mechanically-gated ion channels, recent mechanistic studies in worms have shown that TMC-1 and TMC-2 regulate membrane excitability and egg-laying behavior by conferring a leak conductance (Yue et al., 2018). This raises the question of whether mammalian TMC1 and 2 only function as components of mechanically-gated ion channels or play additional roles that are critical for mechanosensory hair cell function.

In this study, we therefore set out to determine the non-MET functions of TMCs and to tackle its link with hair-cell function, by applying approaches to manipulate TMCs and monitor membrane excitability in mouse hair cells. We seek potential molecular and cellular mechanisms underlying TMCs and the correlated relevance in auditory transduction.

## RESULTS

### TMC1 but not TMC2 mediates a background current in hair cells

During whole-cell patch-clamp recordings from hair cells (Figure 1A, mostly P6 apical-middle OHCs if not specified otherwise) in regular Na^+^-containing external solution (144 mM), we always recorded a significant membrane current (I_m_, 72.69 pA on average) (Figure 1B,C, left). When Na^+^ was replaced in the external solution by N-methyl-D-glucamine (NMDG^+^) (144 mM), the I_m_ was undetectable (Figure 1B, left), demonstrating that this significant background current is carried by an ion channel in the cell membrane. When reperfused with regular external solution, the current baseline returned to “leaky” status (Figure 1B, left). However, the I_m_ was markedly diminished in *Tmc1*-knockout OHCs (Figure 1B,C, right). For more accurate quantification, the amplitude of the background current (I_BG_) was calculated by subtracting the current baseline in NMDG^+^ solution from that in Na^+^ solution (Figure 1D). On average, the I_BG_ in wild-type OHCs was 70.53 pA, while it was drastically reduced to 17.84 pA in *Tmc1*-knockout OHCs (Figure 1D). A more detailed analysis of the I_BG_ was carried out by applying a series of pulse stimulations to hair cells (Figure 1E-G). The IV curves obtained from these measurements verified a greatly diminished membrane current in *Tmc1*-knockout OHCs (Figure 1E,F). By subtraction, it was clear that the inward I_BG_ was dramatically reduced in *Tmc1*-knockout OHCs (Figure 1G).

**Figure 1.**
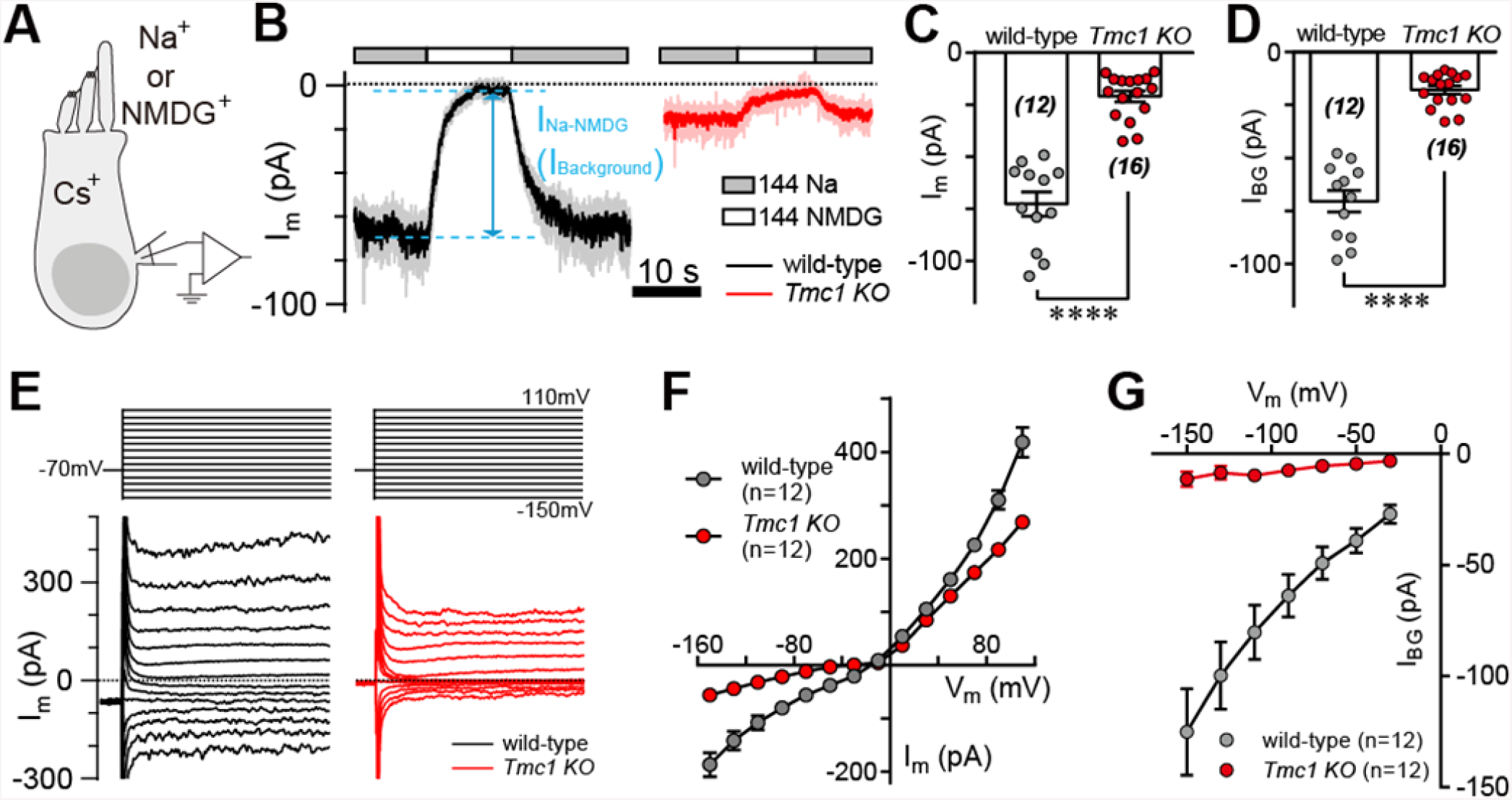
TMC1 mediates a background current in hair cells. (A) Diagram of the recording configuration. The P6 outer hair cells (OHCs) were whole-cell patch-clamped with Cs^+^ in the recording electrode and perfused with different external solutions. (B) Representative traces of membrane current (I_m_) in OHCs from wild-type and *Tmc1*-knockout (*Tmc1 KO*) mice. 144 Na, regular recording solution; 144 NMDG, Na^+^ substituted with NMDG^+^. (C and D) Statistics of the I_m_ (C) and I_Background_ (I_BG_, D) from recordings similar to (B). I_BG_ was calculated by subtraction of I_m_ in 144 Na and 144 NMDG, as indicated in (B). Wild-type I_m_, –72.69 ± 5.79 pA, *Tmc1*-knockout I_m_, –20.96 ± 2.70 pA; wild-type I_BG_, –70.53 ± 5.11 pA, *Tmc1*-knockout I_BG_, –17.84 ± 1.98 pA. (E) Example of I_m_ in OHCs undergoing a series of membrane depolarizations from wild-type (black) and *Tmc1*-knockout (red) mice. (F and G) Statistics of I_m_ (F) and I_BG_ (G) of I-V recordings similar to (E). The external solution contained 1.3 mM Ca^2+^. The holding potential was −70 mV. Data are presented as mean ± SEM. *p <0.05, **p <0.01, ***p <0.001, Student’s t-test.

We next considered whether overexpression of TMC1 would enhance the background current in wild-type hair cells. Three constructs were used for these experiments: enhanced green fluorescent protein control (EGFP), wild-type TMC1 (Tmc1_WT), and TMC1 deafness (Tmc1_dn) carrying a deletion mutation linked to deafness. Using cochlear injectoporation (Xiong et al., 2014), these constructs were delivered into OHCs on postnatal day 3 (P3). The cells were cultured for 1 day *in vitro* (1DIV) and then analyzed by immunostaining (Figure 2A) and patch-clamp recording (Figure 2B). As revealed by HA antibody, exogenously expressed TMC1 largely distributed in soma of OHCs (Figure 2A), consistent with previous observation (Kawashima et al., 2011). While overexpression of the EGFP control and Tmc1_dn did not affect the I_BG_ (17.56 pA and 15.59 pA) (Figure 2C,D), the I_BG_ in OHCs overexpressing Tmc1_WT was increased nearly 2.5 fold (42.53 pA) (Figure 2C,D). These data indicated that hair cells possess a background leak conductance, conferred specifically by TMC1.

**Figure 2.**
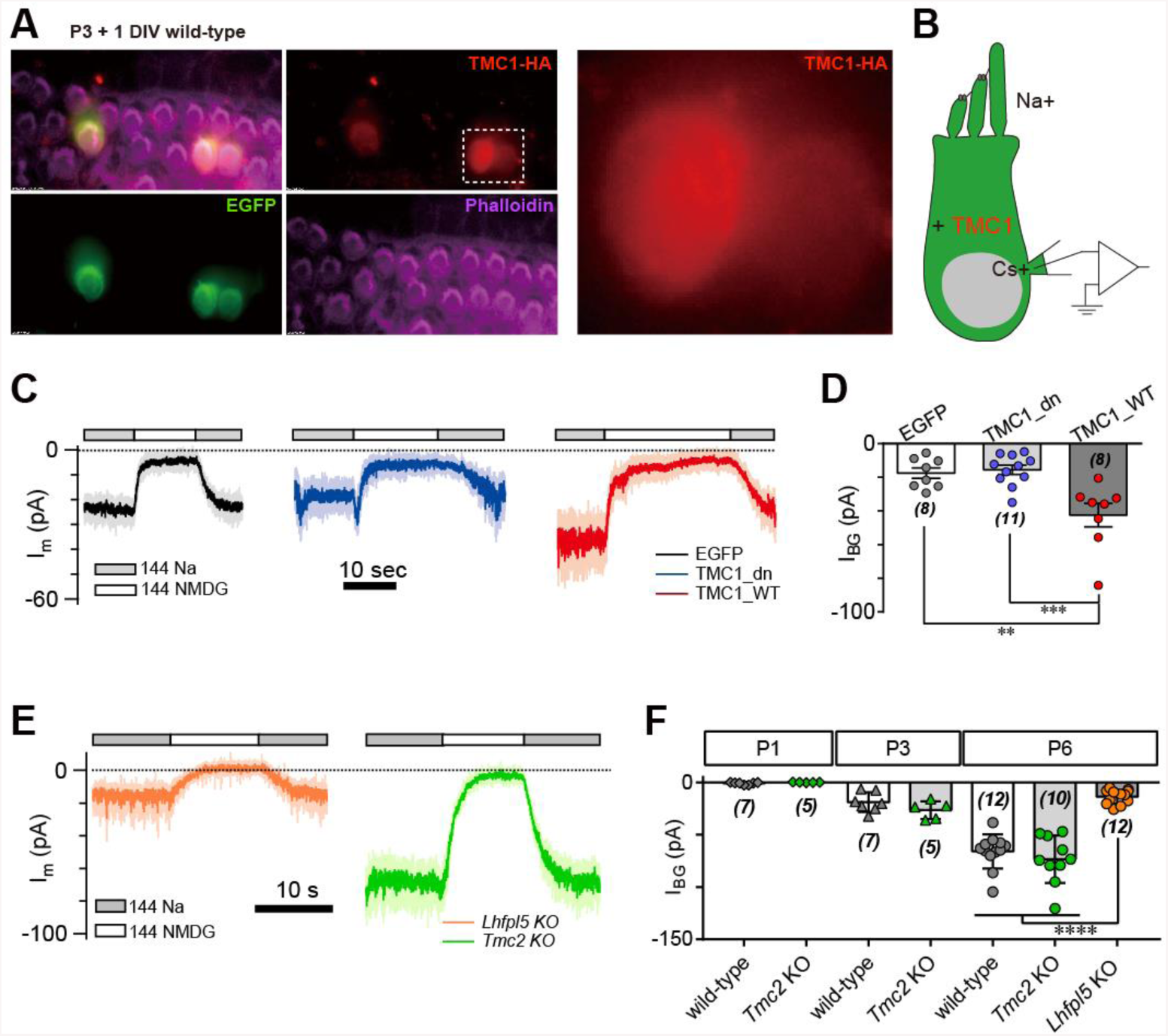
TMC1 but not TMC2 conducts the background current. (A) Exogenous expression of TMC1 in wild-type OHCs from organotypic P3 cochlear tissue cultured for 1 day *in vitro* (P3 + 1DIV). EGFP was co-expressed as an indicator. The OHCs were stained to show spatial distribution of TMC1 (recognized by HA antibody, red), EGFP (by antibody, green), and actin enriched stereocilia (by Phalloidin, magenta), with two OHCs in white dashed frame shown in details. (B) Diagram of the recording configuration. The OHCs were expressing engineered TMC1 with EGFP and whole-cell patch-clamped with Cs^+^ in the recording electrode and Na^+^ extracellularly. (C) Examples of I_m_ of wild-type OHCs at P3 + 1DIV, expressing control (EGFP), deafness TMC1 (TMC1_dn), or wild-type TMC1 (TMC1_WT). (D) Statistical data of I_BG_ from wild-type OHCs expressing EGFP, TMC1_dn, and TMC1_WT under conditions similar to those in (C). I_BG_ values: EGFP, –17.56 ± 3.15 pA; TMC1_dn, –15.59 ± 2.79 pA; TMC1_WT, –42.53 ± 6.96 pA. (E) Representative traces of I_BG_ in cultured OHCs (P6 + 1 DIV) from *Tmc2*- and *Lhfpl5*-knockout mice. (F) Statistics of I_BG_ of OHCs from P2, P4, and P6 *Tmc2*- and *Lhfpl5*-knockout mice. Recordings were made under conditions similar to those in (E). I_BG_ values: P1 wild-type, –0.93 ± 0.40 pA, P1 *Tmc2*-knockout, –0.006 ± 0.119 pA; P3 wild-type, –18.85 ± 3.53 pA, P3 *Tmc2*-knockout, –26.45 ± 3.84 pA; P6 wild-type, –65.94 ± 4.74 pA, P6-knockout, –73.31 ± 7.17 pA, P6 *Lhfpl5*-knockout, –13.64 ± 1.96 pA. The external solution contained 1.3 mM Ca^2+^. The holding potential was −70 mV. Data are presented as mean ± SEM. *p <0.05, **p <0.01, ***p <0.001, Student’s t-test.

It has been suggested that TMC2 is closely coupled with TMC1 in MET function. *Tmc2* expression in the cochlea is highest between P1 and P3, then falls from P4 (Kawashima et al., 2011). Exogenously expressed TMC2 was significantly located in hair bundle of OHCs, as shown by HA tag (Figure S1). We further examined the extent to which TMC2 could confer a background current. Our data showed that the I_BG_ was not altered in *Tmc2*-knockout OHCs at P1, P3, and P6 compared to controls (Figure 2E,F). Similarly, overexpression of TMC2 did not noticeably change the I_m_ baseline (data not shown). In parallel, we analyzed the I_BG_ in *Lhfpl5*-knockout OHCs. Interestingly, similar to *Tmc1*-knockout, there was no evident I_BG_ in *Lhfpl5*-knockout OHCs (Figure 2E,F), consistent with the previous findings that LHFPL5 function in a common pathway (Beurg et al., 2015b; Xiong et al., 2012).

### TMC1-mediated leak current is not carried by the resting open MET channel

Due to existing tension of the hair bundle, there is a sustained open probability of MET channels in hair cells at rest (Assad and Corey, 1992; Corey and Hudspeth, 1983; Johnson et al., 2012). The background current may be composed of the leak current and the resting MET current. To determine the relationship between those current, we therefore analyzed the leak current during mechanical stimulation of hair bundles and in the presence of MET channel blockers (Figure 3A). Since conductance through the MET channel is enhanced when the external Ca^2+^ concentration is low, we carried out the experiments in 0.3 mM Ca^2+^ to increase the sensitivity. At rest, the I_m_ was 97.5 pA (Figure 3A). A sinusoidal mechanical stimulation delivered by a fluid jet deflected hair bundles back and forth to open and close MET channels (Figure 3A, inset). We recorded typical MET current at open status, while the I_m_ at closed status was around 45 pA (Figure 3B, #1, left), similar to the I_m_ as OHCs treated with dihydrostreptomycin (DHS) (Figure 3B, #2, middle). When switching 144 Na^+^ + 100 µM DHS to 144 mM NMDG^+^ solution, the current baseline was near zero (Figure 3B, #3, right), which was considered as I_Leak_ likely mediated by TMC1. Thus, neither mechanical nor pharmacological blockade of the MET channel affect I_Leak_ (Figure 3C). We further examined the proportion of I_Leak_ in I_BG_ in different Ca^2+^ concentration, which became a major part when [Ca^2+^]_o_ was 1.3 mM and larger (Figure 3D). In the following experiments, we presented most of data as I_Leak_ in 1.3 mM [Ca^2+^]_o_ by subtraction current with or without 100 µM DHS.

**Figure 3.**
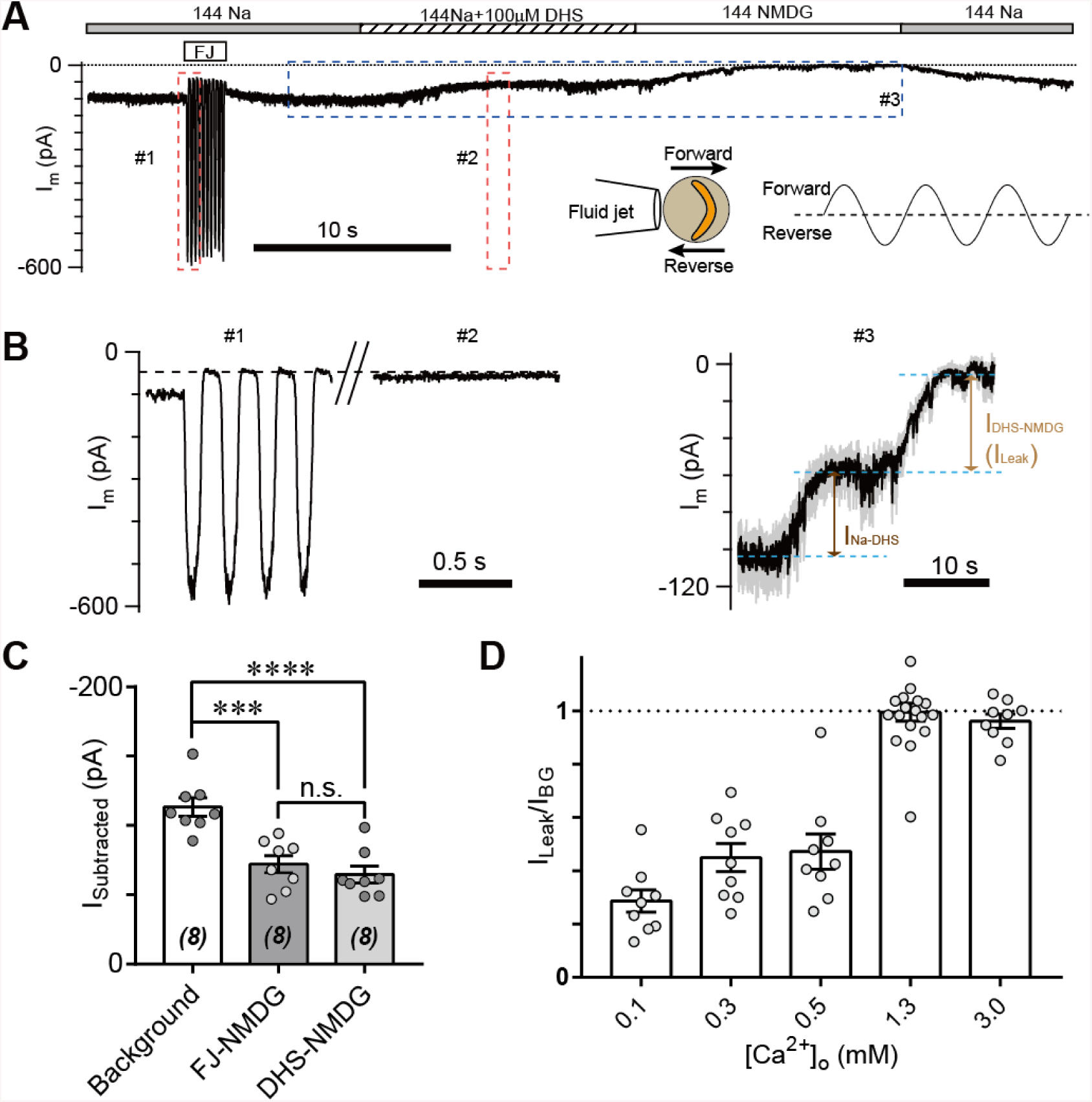
TMC1-mediated leak current is not carried by the resting open MET channel. (A) Representative I_m_ trace showing fluid jet (FJ)-induced open and closed status of MET current and DHS-induced alteration of baseline current. The OHCs were bathed in external solution with 0.3 mM Ca^2+^ instead of 1.3 mM Ca^2+^. Insets: left, a diagram of fluid jet stimulation on a hair bundle; right, a 40 Hz sinusoidal stimulation protocol was used to induce forward and reverse deflection of the hair bundle. (B) Dashed frames #1, #2, and #3 in (A) were shown as enlarged traces. The baseline current was similar when the MET channels were closed by either FJ (#1) or DHS (#2), as highlighted with a dashed line. As shown in #3, the DHS-sensitive resting MET current (I_Na-DHS_) was calculated by subtraction of I_m_ in 144 Na and 144 Na + 100 µM DHS. The baseline current was further closed by NMDG, defined as I_Leak_ by subtraction of I_m_ in 144 Na + 100 µM DHS and 144 NMDG. (C) Statistics of subtracted currents under different conditions: Background, –113.3 ± 6.6 pA; FJ-NMDG (I_Leak_ subtracted from current baseline closed at negative FJ), –72.08 ± 6.18 pA; DHS-NMDG (I_Leak_ subtracted from that closed by 100 µM DHS), –64.63 ± 5.96 pA. (D) Statistics of ratio of I_Leak_ to I_BG_ (I_Leak_ /I_BG_) under different [Ca^2+^]_o_ conditions: 0.1 mM, 0.29 ± 0.04; 0.3 mM, 0.45 ± 0.05; 0.5 mM, 0.47 ± 0.07; 1.3 mM, 1.00 ± 0.03; 3.0 mM, 0.96 ± 0.03. The external solution contained variable Ca^2+^ concentration as indicated. The holding potential was −70 mV. Data are represented as mean ± SEM. *p <0.05, **p <0.01, ***p <0.001, Student’s t-test.

### Amino-acid substitutions in TMC1 alter the TMC1-mediated leak current

We next addressed whether TMC1 itself carries the leak current or is associated with another channel that carries the current. It has been reported that six amino-acids in TMC1 are critical for MET channel function by affecting the pore properties of the channel (Pan et al., 2018) (Figure 4A). We replaced these six amino-acids with cysteine, as reported by Pan et al. (2018), and expressed the mutations in *Tmc1*-knockout OHCs by injectoporation to assess the effects on the leak current (Figure 4B). As controls, we used TMC1_WT and TMC1_dn, and found that the I_Leak_ in *Tmc1*-knockout OHCs at P3+1DIV was restored by TMC1_WT but not by TMC1_dn (Figure 4C,D). Among the cysteine-substituted TMC1 constructs, 5 out of the 6 amino-acids failed to restore the leak current. Especially the G411C, N447C, D528C, and D569C mutations nearly abolished I_Leak_, while T532C partially restored it. Surprisingly, M412C, which has been linked to deafness in *Beethoven* mice (Vreugde et al., 2002), behaved like wild-type TMC1.

**Figure 4.**
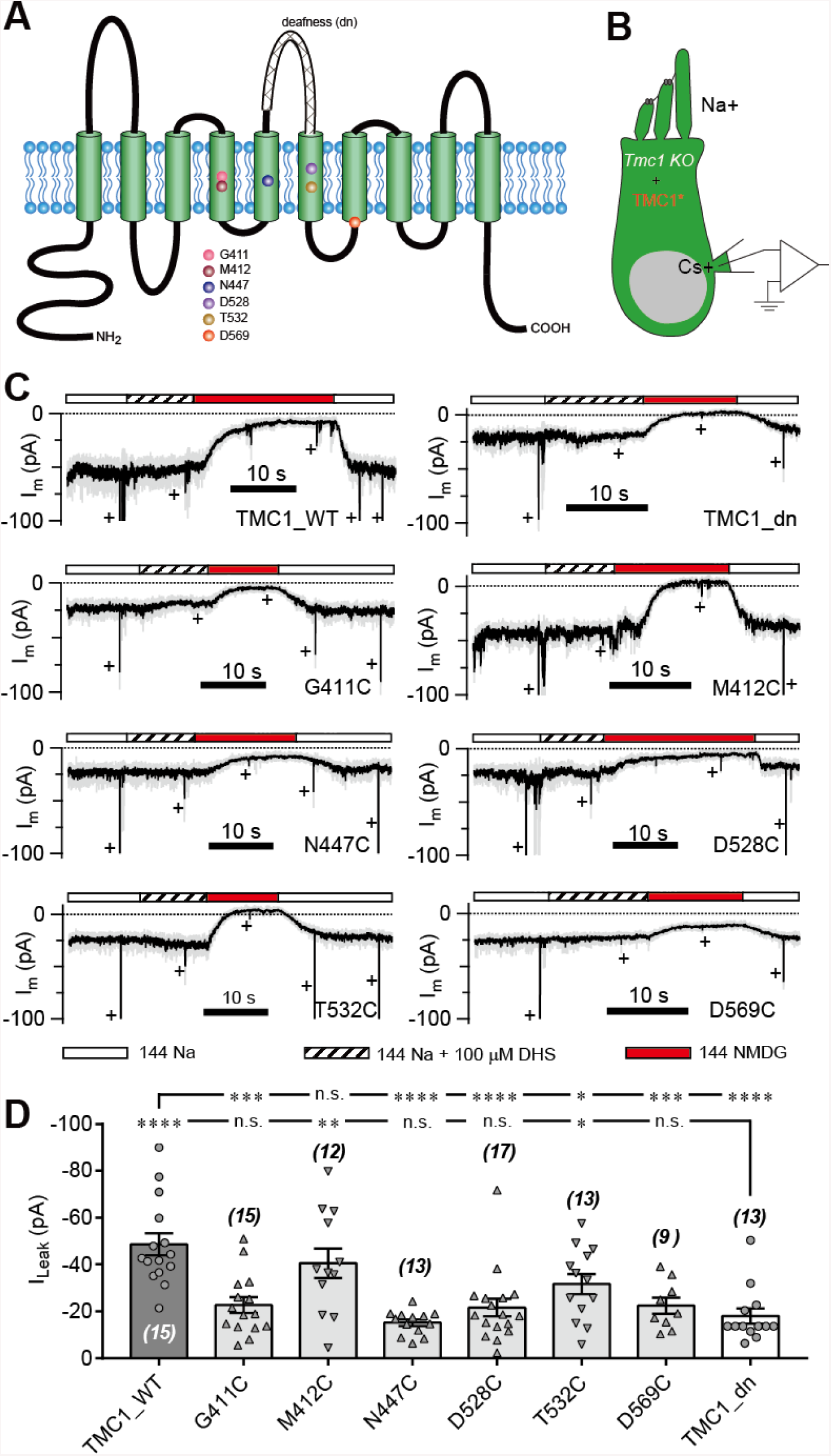
Amino-acid substitution in TMC1 alters the leak current. (A) TMC1 with 10 putative transmembrane domains. The 6 substituted amino-acids are highlighted as colored balls in the predicted positions, and the deafness truncation is at the third extracellular loop between TM5 and TM6. (B) Diagram of the analysis of leak current in cultured *Tmc1*-knockout OHCs (P3 + 1 DIV) expressing modified TMC1 (TMC1*). (C) Representative traces showing the rescue of leak conductance in OHCs by control full-length TMC1 (FL), deafness TMC1 (dn), TMC1-G411C (G411C), TMC1-M412C (M412C), TMC1-N447C (N447C), TMC1-D528C (D528C), TMC1-T532C (T532C), and TMC1-D569C (D569C). Perfusion contents are indicated below. An 800 nm step deflection was applied to the hair bundle by a glass probe. The glass probe induced MET currents are marked “+”, accompanying artefacts induced by switching the perfusion system. Note the MET current was truncated for better showing the leak current. (D) Statistics of rescue by mTMC1 constructs. I_Leak_ values: FL, –48.66 ± 4.76 pA, G411C, –22.69 ± 3.37 pA; M412C, –40.6 ± 6.4 pA, N447C, –15.24 ± 1.42 pA; D528C, –21.66 ± 3.78 pA, T532C, –31.73 ± 4.32 pA, D569C, –22.51 ± 3.43 pA, dn, –18.05 ± 3.25 pA. The rescue indexes of FL and dn were used to evaluate significant difference. Cell numbers were shown on each bar. The external solution contained 1.3 mM Ca^2+^. The holding potential was −70 mV. Data are presented as mean ± SEM. *p <0.05, **p <0.01, ***p <0.001, Student’s t-test.

Treatment with MTSET (2-(trimethylammonium)ethyl methanethiosulfonate, bromide) did not change the current baseline in OHCs expressing any of the six cysteine-substituted TMC1 constructs (Figure S2A). This was not because of the insensitivity of cysteine or a weak MTSET effect, since MTSET treatment did change the MET current amplitude in *Tmc1;Tmc2* double-knockout OHCs expressing M412C (Figure S2B) as previously reported (Pan et al., 2018). The cysteine replacement did not show a consistent pattern of modulation of the leak current and the MET current as summarized in Fig. S2C, implying different molecular mechanisms underlying the two types of current.

### Pharmacological blockade of the TMC1-mediated leak conductance

Next, we set out to evaluate the properties of the leak current by further analyzing its response to pharmacological inhibitors of the MET channel. We first examined the inhibitory effects of the commonly-used MET channel blockers DHS, d-tubocurarine (dTC), and amiloride (Figure 5A-D). DHS had no blocking effect on the background current baseline at a working concentration (100 µM) that blocks MET channels (Figure 5A,B). However, the background leak conductance was 50% inhibited at 487 µM DHS from the fit, 30-times the IC_50_ of the MET channel (Figure 5A,B). dTC and amiloride also affected the leak current, albeit at higher concentrations than the MET current (Figure 5C,D).

**Figure 5.**
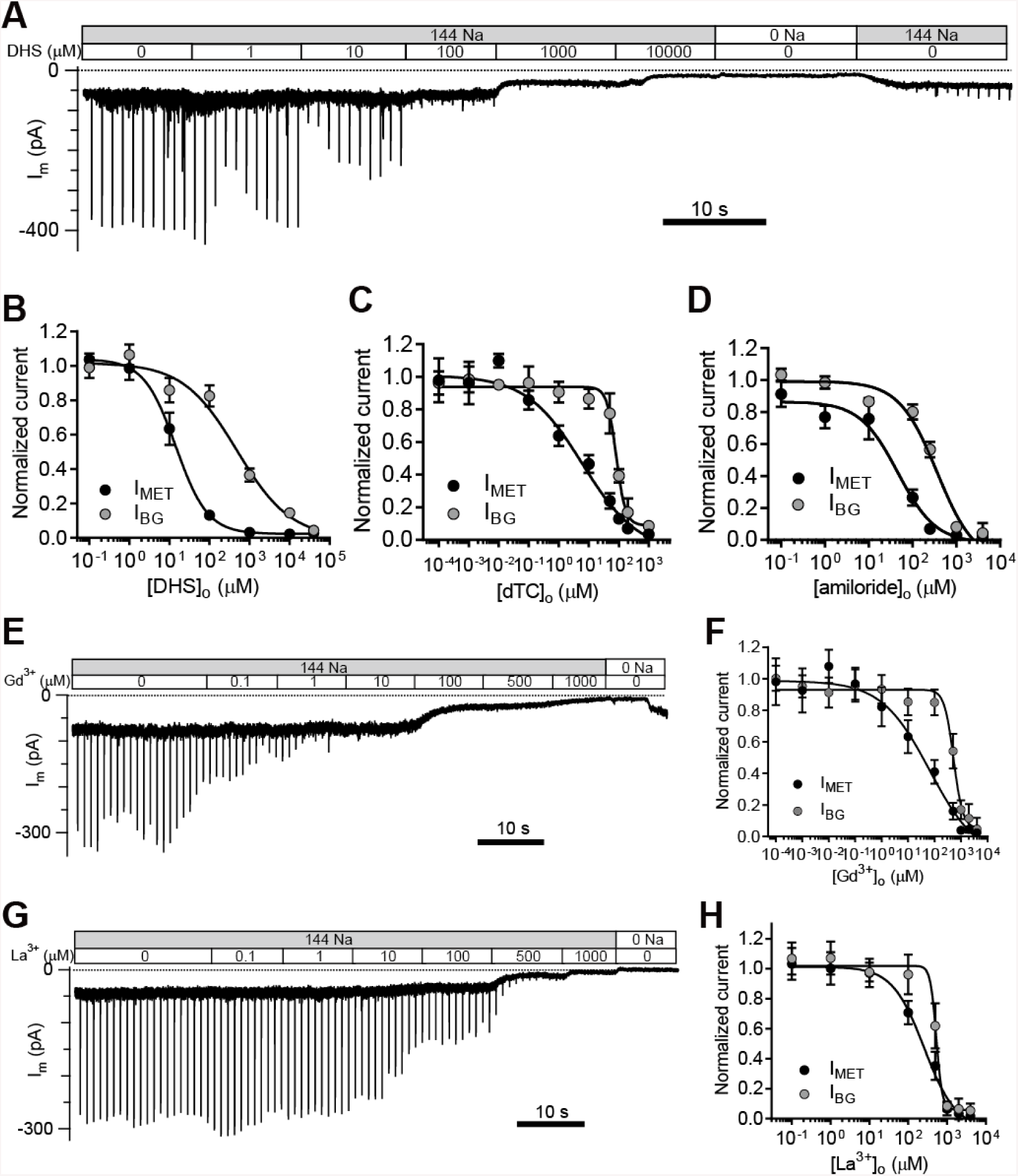
TMC1-mediated leak conductance is antagonized by MET channel blockers. (A and B) Representative trace (A) and statistical curve (B) of I_m_ inhibition by DHS. A train of 800 nm step deflection was applied to the hair bundle by a glass probe to induce MET currents. I_MET_ and I_BG_ were calculated and plotted against the DHS concentration. As fitted by, the IC_50_ of DHS was 15 µM for the MET channels and 487 µM for the leak conductance (cell numbers, 7–11). Hill slope: I_MET_, −1.10; I_BG_, −0.65. (C and D) Statistical dose curve of I_m_ with graded concentrations of d-tubocurarine (dTC) (C) and amiloride (D). dTC IC_50_: I_MET_, 6 µM; I_BG_, 82 µM. dTC Hill slope: I_MET_, −0.47; I_BG_, −2.80. dTC cell numbers, 5–15. Amiloride IC_50_: I_MET_, 46 µM; I_BG_, 365 µM. Amiloride Hill slope: I_MET_, −1.36; I_BG_, −1.67. Amiloride cell numbers, 7–16. (E and F) Dosage effect of Gd^3+^. Example trace (E) and statistical curve (F) of I_m_ in OHCs during perfusion with solutions containing graded concentrations of Gd^3+^. A train of 800 nm step deflection was applied to the hair bundle by a glass probe to induce MET currents. The MET and leak current amplitudes changed due to the channel sensitivity of Gd^3+^ and NMDG. IC_50_: I_MET_, 66 µM; I_BG_, 524 µM. Hill slope: I_MET_, −0.48; I_BG_, −2.49. Cell numbers, 7–16. (G and H) Dose effect of La^3+^. Example trace (G) and dosage curve (H) of I_m_ with La^3+^ treatment. A train of 800 nm step deflection was applied to the hair bundle by a glass probe to induce MET currents. IC_50_: I_MET_, 259 µM; I_BG_, 531 µM. Hill slope: I_MET_, −1.06; I_BG_, −5.67. Cell numbers, 7–8. For space reason, 144 NMDG was shown as 0 Na. The external solution contained 1.3 mM Ca^2+^. The holding potential was −70 mV. Data are presented as mean ± SEM. *p <0.05, **p <0.01, ***p <0.001, Student’s t-test.

It has been reported that trivalent cations, such as Gd^3+^ and La^3+^, block MET channels (Farris et al., 2004; Kimitsuki et al., 1996), so we applied Gd^3+^ and La^3+^ at various concentrations and monitored the inhibitory effects on evoked MET current and leak current (Figure 5E-H). Surprisingly, the leak current was not affected even when [Gd^3+^]_o_ reached 80 µM, the IC_50_ for blocking the MET current (Figure 5E,F). However, the leak current was inhibited by [Gd^3+^]_o_ with an IC_50_ of 541 µM (Figure 5E,F). Similarly, [La^3+^]_o_ inhibited the MET channel with an IC_50_ of 259 µM and the leak current with an IC_50_ of 531 µM (Figure 5G,H). Note I_BG_ was shown in Figure 5 but it mostly represented the TMC1 mediated leak component indeed.

### Ionic permeability of the TMC1-mediated leak conductance

To further characterize the leak current in OHCs, we carried out a series of ion-permeation tests using the cations Li^+^, Cs^+^, Ba^2+^, Zn^2+^, Co^2+^, Mg^2+^, and Ca^2+^ (Figure 6A,B). Most of the cations shared a size of I_Leak_ similar to Na^+^, except for Cs^+^ and Ca^2+^ (Figure 6A,B). The Cs^+^-conducted I_Leak_ was slightly larger (Figure 2A), while 75 mM Ca^2+^ robustly blocked the I_Leak_ (Figure 6B). The Ca^2+^ permeability of the leak channel was further determined from calculation of reversal potentials by a voltage ramp stimulation with Ca^2+^ extracellularly and Cs^+^ intracellularly (Figure 6C). The Ca^2+^ permeability was extremely small comparing to Na^+^ and Mg^2+^ permeability (Figure 6D). Next, we monitored the background and MET currents in solutions containing different concentrations of Ca^2+^ and Na^+^. Significantly, the background current was highly sensitive to Ca^2+^; it increased when [Ca^2+^]_o_ declined and decreased when [Ca^2+^]_o_ increased, while the MET current was reduced at first and then reached a plateau after [Ca^2+^]_o_ was >10 mM (Fig. 6E,F) that was sufficient to block the membrane current to an extent similar to TMC1 removal in OHCs (Fig. 6G).

**Figure 6.**
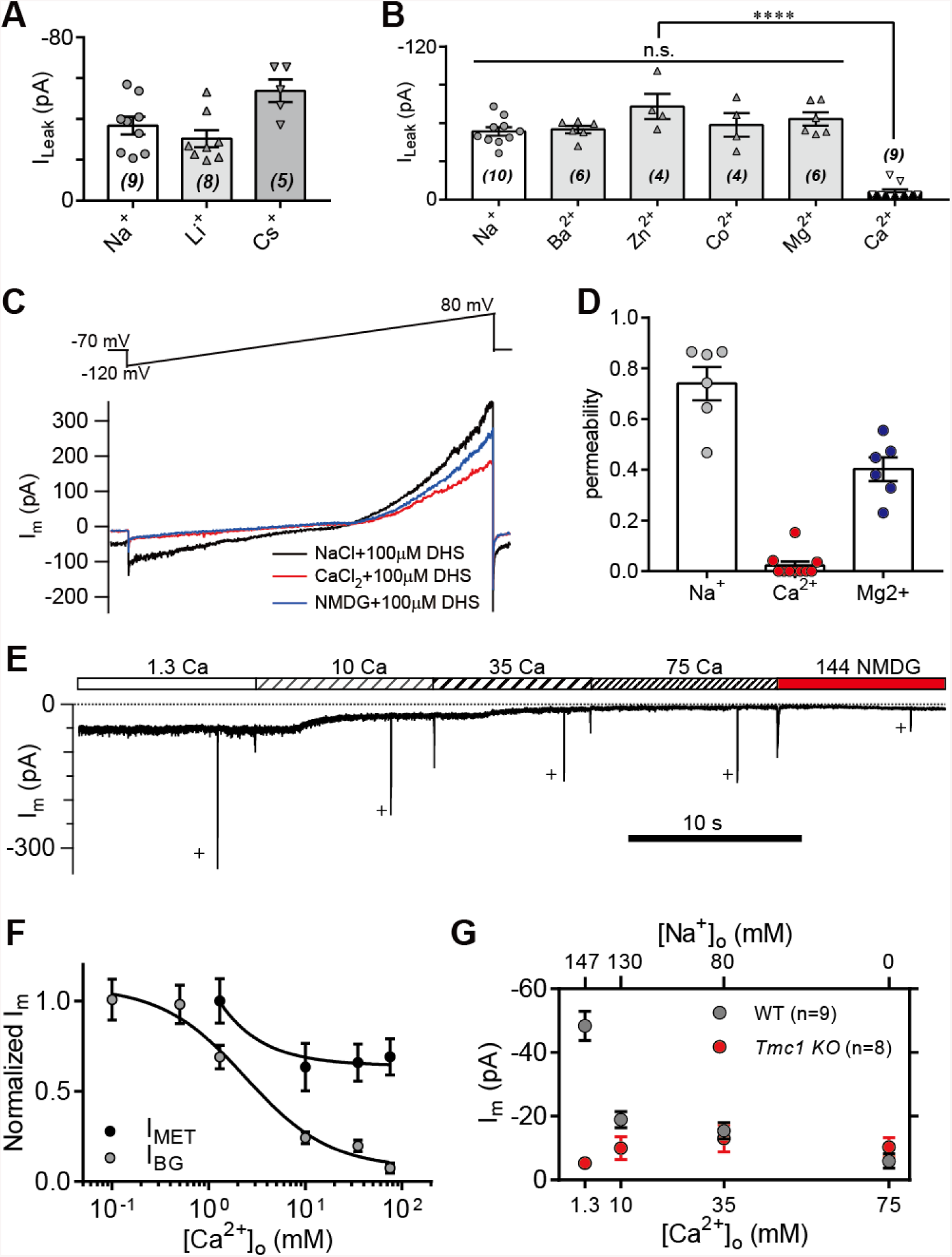
High-concentration Ca^2+^ blocks the leak current but not MET current. (A) Monovalent cations Li^+^ and Cs^+^ conducted the leak current. In this experiment, 150 mM NaCl was substituted with 150 mM LiCl or 150 mM CsCl in the external solution. (B) Divalent cations 10 mM Ba^2+^, 75 mM Zn^2+^, 75 mM Co^2+^, 150 mM Mg^2+^, and 75 mM Ca^2+^, conducted the leak current. The 150 mM NaCl was partially or completely replaced with the according cations as described in the Methods. (C) Representative I_m_ traces by ramp stimulation for calculation of ionic permeability. Extracellular ion was switched from 150 mM Na^+^ to 75 mM Ca^2+^ + 75 mM NMDG^+^, and to 150 NMDG^+^. In the intracellular solution, 150 mM CsCl was used. (D) Statistics of ionic permeability calculated from similar recordings in (C). (E) Example trace of I_m_ of OHCs during perfusion with solutions containing graded concentrations of Ca^2+^ and Na^+^. An 800 nm step deflection was applied to the hair bundle by a glass probe. The glass probe induced MET currents are marked “+”, accompanying artefacts induced by switching the perfusion system. (F) Dose curves of I_BG_ and I_MET_ in wild-type OHCs in different Ca^2+^ and Na^+^ concentrations (cell numbers, 9–20). (G) Statistical analysis of dose-dependent background leak current in OHCs from wild-type (black) and *Tmc1*-knockout (red) mice when bathed in mixed Ca^2+^ and Na^+^. The ions and concentrations used in test external solutions were variable, as described in this figure legend and the methods. The holding potential was −70 mV. Data are presented as mean ± SEM. *p <0.05, **p <0.01, ***p <0.001, Student’s t-test.

### The leak current modulates action potential firing in IHCs

We next tackled the physiological relevance of the TMC1-mediated background conductance in auditory transduction. A significant leak conductance would be expected to depolarize the membrane potential and affect cell excitability. IHCs are innervated by the spiral ganglion neurons that transmit sound information to the CNS and signal transmission from hair cells to the spiral ganglion might therefore be affected by the leak conductance. We therefore measured the membrane potential (V_m_) in IHCs (Figure 7A). In wild-type IHCs, the V_m_ varied actively and periodically in the bursting and non-bursting states (Figure 7A). However, the V_m_ was largely hyperpolarized and there was almost no action potential firing in *Tmc1*-knockout IHCs (Figure 7A). With positive current injection, the *Tmc1*-knockout IHCs fired action potentials at threshold similar to wild-type IHCs (Figure 7A). Although the V_m_ in the non-bursting state was more hyperpolarized than in the bursting state in wild-type IHCs, it was positive to the V_m_ in *Tmc1*-knockout IHCs (Figure 7A,B). This change of membrane excitability was also defined by monitoring the action potential bursting rate (Figure 7C) and the leak current (Figure 7D). The leak current was smaller in IHCs than that in OHCs, which may be due to different expression profile of potassium channels (Marcotti et al., 2006; Marcotti et al., 2003). With ramp current injection, the firing threshold was similar, but the minimum injected current required to induce firing in *Tmc1*-knockout IHCs was ∼20 pA greater than that in wild-type IHCs (Figure 7E,G,H). When depolarized by stepped current injection, the firing rate was lower in *Tmc1*-knockout IHCs and the rate-current curve was shifted to the right but finally reached a similar level when a larger current was injected (Figure 7F,I).

**Figure 7.**
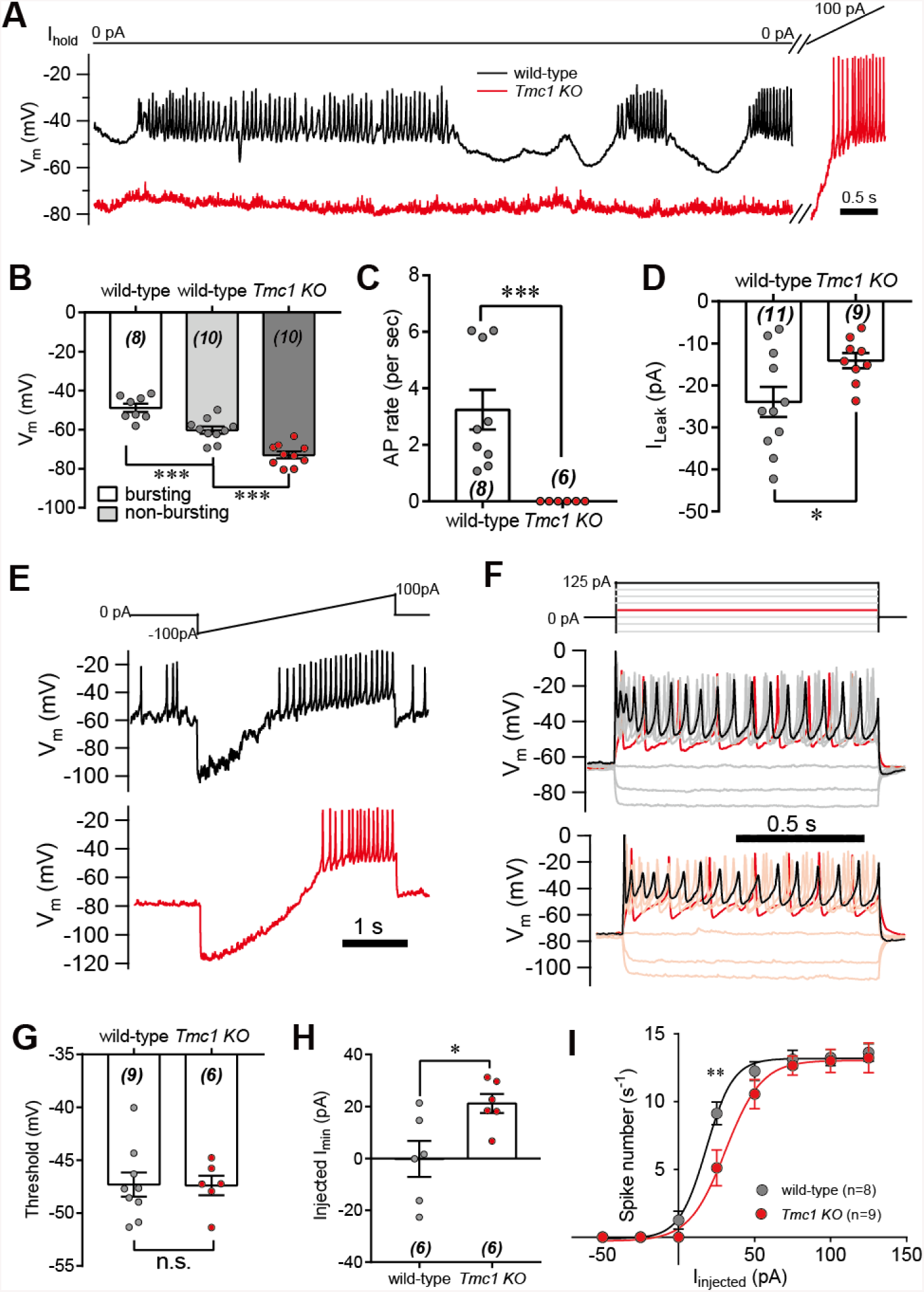
IHC excitability is down-regulated in *Tmc1*-knockout mice. (A) Representative current-clamp recording in IHCs from wild-type (black) and *Tmc1*-knockout (red) mice. For the most part, the IHCs were held at 0 pA. To define the excitability, a ramp current was injected into the *Tmc1*-knockout IHCs to induce a burst of spikes. (B) Statistics of V_m_ recorded in IHCs similar to (A). Values of V_m_ in wild-type IHCs were defined as two states, bursting and non-bursting, which did not apply to *Tmc1*-knockout IHCs. V_m_ of wild-type in bursting state, 48.78 ± 2.15 mV; wild-type in non-bursting state, 60.19 ± 1.86 mV; *Tmc1*-knockout, 72.89 ± 1.75 mV. (C) Statistics of firing rate (spikes/s) in IHCs similar to (A). Values of firing rate: wild-type, 3.2 ± 0.7 Hz; *Tmc1*-knockout, 0 ± 0 Hz. (D) Statistics of I_Leak_ from voltage-clamp recording in IHCs. Values of I_Leak_: wild-type, 23.93 ± 3.58 pA; *Tmc1*-knockout, 14.1 ± 1.79 pA.(E) Representative current-clamp traces of V_m_ in IHCs with ramp-current injection from –100 pA to +100 pA for 3 s. (F) Representative current-clamp recording in IHCs stimulated by a family of depolarization currents from –50 pA to +125 pA at 25 pA steps. (G) Statistics of firing threshold from data as in (E). Values of threshold were –47.3 ± 1.2 mV in wild-type OHCs and –47.39 ± 0.92 mV in *Tmc1*-knockout OHCs. (H) Statistics of minimum current injected (Injected I_min_) to evoke an action potential from data as in (E). In wild-type OHCs: –0.23 ± 6.95 pA; in *Tmc1*-knockout OHCs: –21.12 ± 3.66 pA. (I) Statistics of numbers of spike/s from data as in (F). wild-type: 0 pA, 1.25 ± 0.67; 25 pA, 9.13 ± 0.83; 50 pA, 12.25 ± 0.70; 75 pA, 13.125 ± 0.67, 100 pA, 13.25 ± 0.59; 125 pA, 13.625 ± 0.65. *Tmc1*-knockout: 0 pA, 0 ± 0; 25 pA, 5.11 ± 1.32; 50 pA, 10.56 ± 1.08; 75 pA, 12.67 ± 0.71, 100 pA, 13.00 ± 0.85; 125 pA, 13.22 ± 1.10. The external solution contained 1.3 mM Ca^2+^. K^+^ was used in the intracellular solution for current-clamp recordings in this figure except that Cs^+^ was used for voltage-clamp recording in (D). Data are presented as mean ± SEM. *p <0.05, **p <0.01, ***p <0.001, Student’s t-test.

### The leak current follows the tonotopic gradient of the MET response in OHCs

Next, we investigated the effect of the leak current in OHCs. First, the I_Leak_ was examined in OHCs along the cochlear coil (Figure 8A). We indeed found a gradient in the leak current in wild-type OHCs, while the gradient was abolished in *Tmc1*-knockout OHCs (Figure 8B,C). Next, we analyzed the MET current along the cochlear coil when blocking the leak current with 35 mM [Ca^2+^]_o_ since 35 mM [Ca^2+^]_o_ was sufficient to block the leak current to an extent similar to TMC1 removal in OHCs (Figure 6G). Strikingly, the gradual increase in MET current amplitude was severely blunted in OHCs in the presence of 35 mM [Ca^2+^]_o_ (Figure 8D,E). The effect was reminiscence to that previously reported (Beurg et al., 2014) and to our observation (Figure 8D,E) for hair cells lacking TMC1. The MET current decreased from apex to base in *Tmc1*-knockout OHCs, which might result from that the hair bundle got more disrupted at base coil. These data suggested that the tonotopic gradient of TMC1-mediated leak current and MET current in OHCs could be modulated by external Ca^2+^.

**Figure 8.**
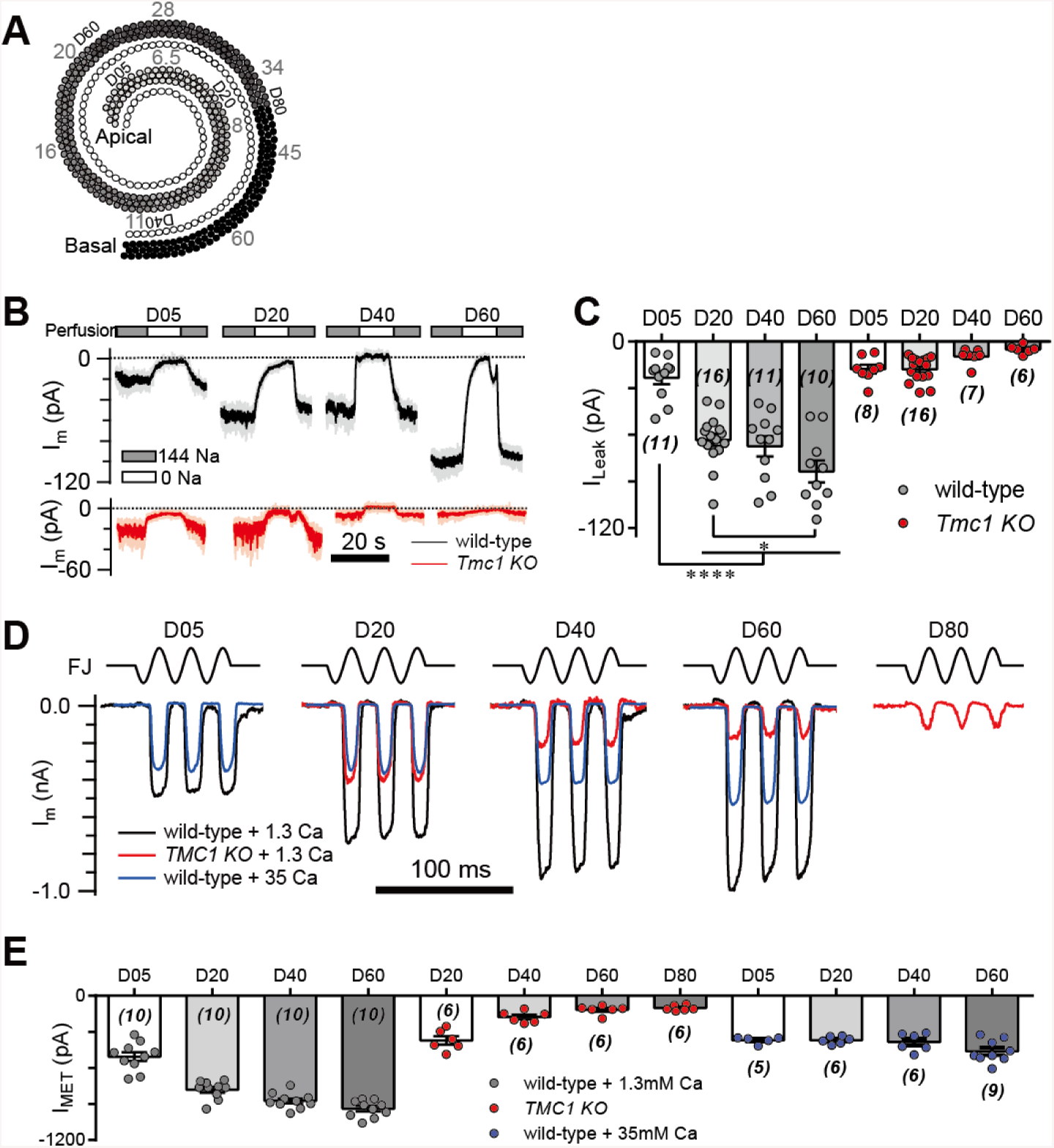
TMC1-mediated leak and MET currents in OHCs. (A) Diagram showing the tonotopic map in mouse hair cells (adapted from Figure 1B in Kim and Fettiplace, 2013), labelled with response frequencies (kHz, grey) and location (D% to apex, black). The apex and base are defined as 0 and 1, with reference to which D05, D20, D40, D60, and D80 represent distances of 0.05, 0.2, 0.4, 0.6, and 0.8. (B) Representative traces of I_m_ recorded in OHCs at different locations along the cochlear coil, from wild-type (black) and *Tmc1*-knockout (red) mice. The external solution contained 1.3 mM Ca^2+^. The apex and base are defined as 0 and 1, with reference to which D05, D20, D40, and D60 represent distances of 0.05, 0.2, 0.4, and 0.6. (C) Statistical analysis of location-specific I_Leak_ from similar recordings as in (B). Values of I_Leak_ in wild-type OHCs (pA): D05, –23.3 ± 4.1; D20, –63.26 ± 3.98; D40, –67.45 ± 6.51; D60, –83.53 ± 7.14. I_Leak_ values in *Tmc1*-knockout OHCs (pA): D05, –17.72 ± 2.82; D20, –17.84 ± 1.98; D40, –9.56 ± 1.87; D60, – 4.98 ± 1.15. (D) Representative traces of location-specific MET current in wild-type OHCs when bathed in 1.3 mM or 35 mM Ca^2+^ and *Tmc1*-knockout OHCs when bathed in 1.3 mM Ca^2+^. A sinusoidal deflection was applied to the hair bundle by a fluid jet. (E) Statistical analysis of location-specific macroscopic MET current. Values of I_MET_ in wild-type OHCs in 1.3 mM Ca^2+^ (pA): D05, –505.1 ± 37.2 pA; D20, –780.2 ± 23.7 pA; D40, –872.3 ± 20.5 pA; D80, –938.8 ± 21.8 pA. Values of I_MET_ in wild-type OHCs in 35 mM Ca^2+^ (pA): D05, –369 ± 12.6 pA; D20, –368.9 ± 126 pA; D40, –384.4 ± 30.2 pA; D60, –461.1 ± 30.6 pA. Values of I_MET_ in *Tmc1*-knockout OHCs in 1.3 mM Ca^2+^ (pA): D20, –371.2 ± 34.7 pA; D40, –176.8 ± 19.1 pA; D60, – 116.9 ± 15.4 pA; D80, –102.4 ± 9.3 pA. The holding potential was −70 mV. Data are presented as mean ± SEM. *p <0.05, **p <0.01, ***p <0.001, Student’s t-test.

We next determined whether the change in macroscopic MET current represents a change in the unitary MET channel conductance and whether the absence of the leak current disrupts the tonotopic gradient of MET conductance. The unitary MET channel analysis showed that 35 mM [Ca^2+^]_o_ reduced the unitary MET channel current to ∼5 pA in both low-frequency and high-frequency OHCs (Figure 9A,B). These data further suggested that the extracellular Ca^2+^ modulates leak conductance and MET channel properties accordingly.

**Figure 9.**
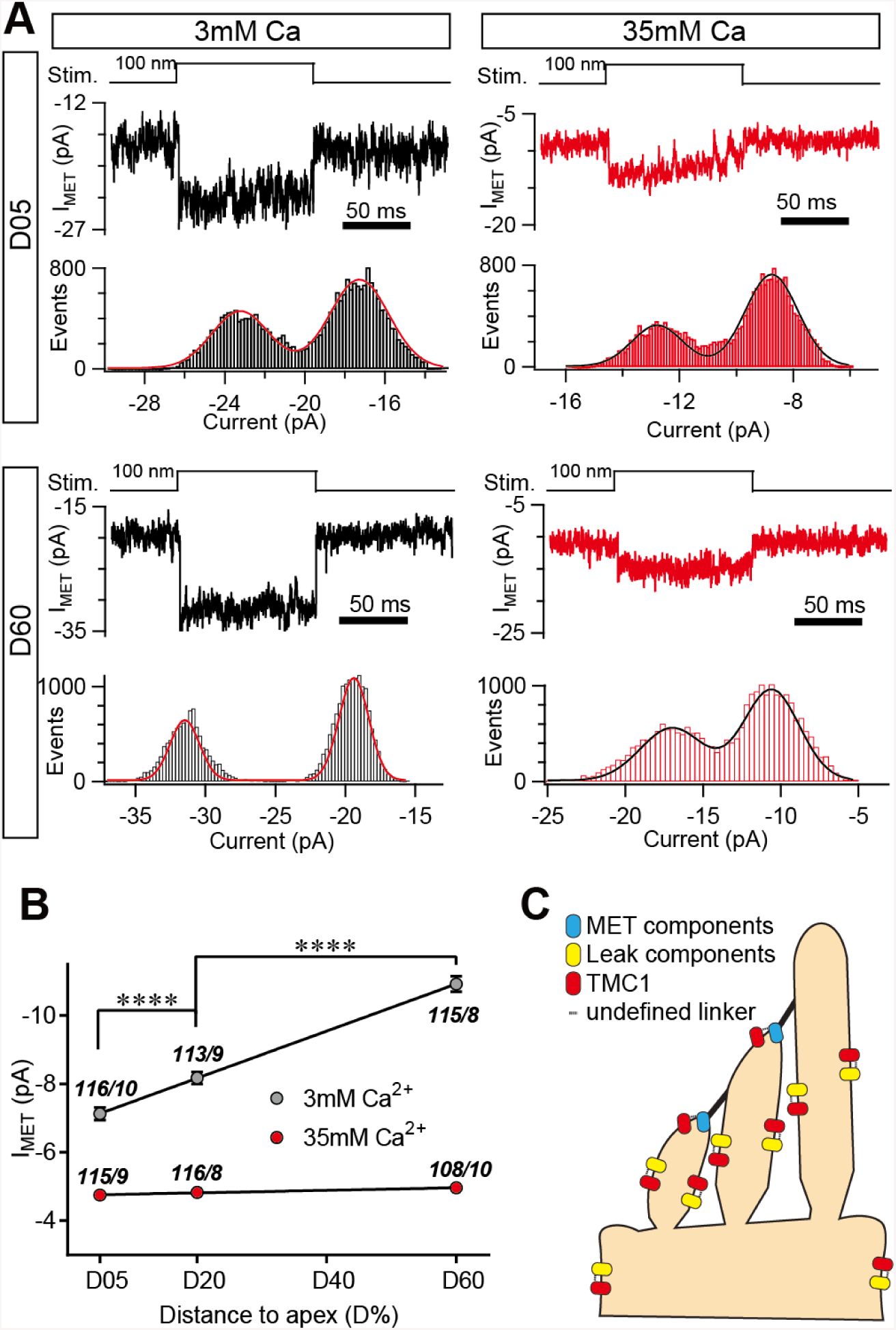
High Ca^2+^ removes the MET conductance gradient as revealed by unitary channel analysis. Location-specific single MET channel recording from wild-type OHCs in solution with 3 mM or 35 mM Ca^2+^ at D05 or D60. The traces were chosen to show nice dual-peak fitting but did not represent normal flickers. A 100-nm step deflection was applied to the hair bundle by a glass probe. Statistical analysis of location-specific unitary MET channel current. Values of unitary I_MET_ in 3 mM Ca^2+^: D05, –7.03 ± 0.19 pA; D20, –7.90 ± 0.16 pA; D60, –10.59 ± 0.19 pA. Values of I_MET_ in 35 mM Ca^2+^: D05, –4.72 ± 0.11 pA; D20, –4.76 ± 0.10 pA; D60, –4.92 ± 0.11 pA. Numbers are shown as events/cells. (C) A working model of the molecular mechanism by which TMC1 tunes both MET and leak channels. Dashed line between TMC1 and other proteins indicates undefined coupling. The holding potential was −70 mV. Data are presented as mean ± SEM. *p <0.05, **p <0.01, ***p <0.001, Student’s t-test.

## DISCUSSION

Here we uncovered, in mammalian hair cells, a previously-unappreciated role of TMC1 in mediating a background conductance and thereby regulating membrane excitability. With TMC1 deficiency, the resting membrane potential is hyperpolarized, resulting in the absence of spontaneous action potential firing in neonatal IHCs (Figure 7) and the removal of gradient of MET conductance in OHCs (Figure 8 and 9) (Beurg et al., 2014; Beurg et al., 2015b). In other species, TMC orthologues function in diverse ways according to their expression pattern in effector cells (Guo et al., 2016; He et al., 2019; Wang et al., 2016; Yue et al., 2018; Zhang et al., 2015; Zhang et al., 2016). It has been recognized that leak conductance is generally used in the nervous system to regulate neuronal excitability and thus circuit activity; it recruits a variety of channels on the plasma membrane or endoplasmic reticulum (Bers, 2014; Enyedi and Czirjak, 2010; Lu et al., 2007). Hence, these results strongly support the hypothesis that the excitability of cells and neural circuits that control processes from sensory transduction to motor function are commonly upregulated by TMC proteins in diverse organisms.

TMC1 is likely a major component of the leak conductance, as implied by the mutagenesis experiment (Figure 4). Our data showed that at least 4 amino-acids are critical for the leak conductance, since these constructs cannot restore the leak current after replacement of a single amino-acid by cysteine, implying that TMC1 is key to generating the leak conductance. However, adding positive charge to these amino-acids does not affect the leak conductance, as revealed by its insensitivity to treatment with MTSET. In addition, the leak conductance is inhibited by typical MET channel blockers, implying that TMC1 as the responsible component. It has been proposed that both TMC-1 and TMC-2 confer the leak conductance in worms (Yue et al., 2018). However, in our study of mice, only TMC1 but not TMC2 mediated the leak conductance (Figure 2), indicating a unique non-MET role of TMC1 in mammals.

Intriguingly, the leak conductance differed from the MET conductance in several properties, although both are functional representations of TMC1. First, the leak current did not stem from the resting open MET channels (Figure 3). Second, the patterns of change differed for the leak conductance and the MET current according to the amino-acid substitution experiment (Figures 4 and S2). Third, the leak channel shared a group of identical antagonists with the MET channel but had different kinetics (Figure 5). Both MET channel blockers (Figure 5A-D) and non-selective cation channel blockers (Figure 5E-H) inhibited the leak current but with an IC_50_ 5–10-fold that for the MET channel. Last, extracellular high Ca^2+^ blocked the leak conductance but not the MET channel (Figure 6). These lines of evidence indicate that TMC1 confers the leak conductance by a mechanism distinct from the MET channel.

Interestingly, the TMC1-mediated leak conductance exhibits a tonotopic pattern in OHCs, in parallel with the tonotopicity of the MET current, which is defined by several lines of evidence. First, we found that the leak current still existed in *Tmc2*-knockout OHCs while it was absent from *Tmc1*- or *Lhfpl5*-knockout OHCs (Figure 2E,F); this is consistent with the finding that the gradient of the MET response was lost in *Tmc1*- and *Lhfpl5*-knockout mice and preserved in *Tmc2*-deficient mice (Beurg et al., 2015b). Second, the leak conductance increased along the cochlear coil (Figure 8A,B), which also coincides with the spatial *Tmc1* expression pattern (Kawashima et al., 2011) and the tonotopic gradient of TMC1 proteins in graded numbers (Beurg et al., 2018). Last, high Ca^2+^ blockade abolished both the background current and the gradient of the MET response, defined by the analysis of the macroscopic (Figure 8) and unitary MET current (Figure 9). The tonotopic gradient of conductance in OHCs is an important property of hair-cell MET (Beurg et al., 2006; Ricci et al., 2003; Waguespack et al., 2007). Our results showed that the background leak conductance, together with the MET response, is tuned by extracellular Ca^2+^ and other unknown determinants, which is not surprising since other factors, such as PIP2, also regulate MET channel pore properties (Effertz et al., 2017).

Due to limited information about the structure of TMC1, we do not yet know how TMC1 confers the leak conductance. It has been shown that only a proportion of TMC1 proteins are localized around tip-links, and the number of TMC1 proteins increases from apex to base in OHCs as reported in a transgenic TMC1 mouse model (Beurg et al., 2018; Kurima et al., 2015), consistent with a scenario in which extra TMC1 proteins that are not in the MET complex provide the leak conductance. Based on current data, we suggest a working model (Figure 9C) in which TMC1 has dual functions by integrating into the MET channel for the mechanically-induced conductance and attaching to other undefined components for the leak conductance, in which the activity of both channels is tuned by TMC1. Interestingly, TMC1 may form dimers by sharing a protein fold similar to TMEM16A, a Ca^2+^-activated Cl^−^ channels or Ca^2+^-activated lipid scramblase (Ballesteros et al., 2018; Kunzelmann et al., 2016; Pan et al., 2018). However, this hypothesis needs to be further tested in structural and functional studies.

## EXPERIMENTAL PROCEDURES

### Mouse strains and animal care

The mouse strains used in this study, B6.129-TMC1<tm1.1Ajg>/J, B6.129-TMC2<tm1.1Ajg>/J, and B6.129-Lhfpl5<tm1Kjn>/Kjn, were from the Jackson Laboratory (Bar Harbor, ME). The experimental procedures on mice were approved by the Institutional Animal Care and Use Committee of Tsinghua University.

### DNA constructs, cochlear culture, and injectoporation

Mouse *Tmc1* and *Tmc2* cDNAs were cloned into CMV-Script and pCDNA3.1-vectors, respectively. To obtain the *Tmc1-deafness* vector and amino-acid-substituted *Tmc1* constructs, specific primers were designed and used for PCR (listed in Table S1). Cochlear culture and injectoporation were as previously described (Xiong et al., 2014). In this study, the organ of Corti was isolated from P3 mice and cut into 3 pieces in Dulbecco’s modified Eagle’s medium/F12 with 1.5 µg/ml ampicillin. For electroporation, a glass pipette (2 µm tip diameter) was used to deliver cDNA plasmids (0.2 µg/µl in 1× Hanks’ balanced salt solution) to hair cells in the sensory epithelium. EGFP was used as an indicator for the selection of transfected hair cells. A series of 3 pulses at 60 V lasting 15 ms at 1-s intervals was applied to cochlear tissues by an electroporator (ECM Gemini X2, BTX, CA). The cochlear tissues were cultured for 1 day *in vitro* and then used for electrophysiological recording.

### Electrophysiology

Hair cells were recorded using whole-cell patch-clamp as previously described (Xiong et al., 2012). All experiments were performed at room temperature (20-25°C). Briefly, the basilar membrane with hair cells was acutely dissected from neonatal mice. The dissection solution contained (in mM): 141.7 NaCl, 5.36 KCl, 0.1 CaCl_2_, 1 MgCl_2_, 0.5 MgSO_4_, 3.4 L-glutamine, 10 glucose, and 10 H-HEPES (pH 7.4). Then the basilar membrane was transferred into a recording chamber with recording solution containing (in mM): 144 NaCl, 0.7 NaH_2_PO_4_, 5.8 KCl, 1.3 CaCl_2_, 0.9 MgCl_2_, 5.6 glucose, and 10 H-HEPES (pH 7.4). For I_leak_ calculation, the cells were further bathed in recording solution containing 144 mM NMDG that replaced 144 mM NaCl. The acutely isolated or cultured basilar membrane was used for electrophysiological recording within 1 h. Hair cells were imaged under an upright microscope (BX51WI, Olympus, Tokyo, Japan) with a 60× water-immersion objective and an sCMOS camera (ORCA Flash4.0, Hamamatsu, Hamamatsu City, Japan) controlled by MicroManager 1.6 software (Edelstein et al., 2010). Patch pipettes were made from borosilicate glass capillaries (BF150-117-10, Sutter Instrument Co., Novato, CA) with a pipette puller (P-2000, Sutter) and polished on a microforge (MF-830, Narishige, Tokyo, Japan) to resistances of 4-6 MΩ. Intracellular solution contained (in mM): 140 CsCl, 1 MgCl_2_, 0.1 EGTA, 2 Mg-ATP, 0.3 Na-GTP, and 10 H-HEPES, pH 7.2), except when CsCl was replaced with KCl in current-clamp. Hair cells were recorded with a patch-clamp amplifier (EPC 10 USB and Patchmaster software, HEKA Elektronik, Lambrecht/Pfalz, Germany). The liquid junction potential is not corrected in the data shown. As measured, the pipette with CsCl intracellular solution had a value of +4 mV in regular recording solution and −6 mV in 144 mM NMDG^+^ solution.

For single-channel recordings, we followed published procedures (Ricci et al., 2003; Xiong et al., 2012). The intracellular solution was the same for macroscopic and microscopic current recording. To break tip-links, hair bundles were exposed to Ca^2+^-free solution using a fluid jet (in mM): 144 NaCl, 0.7 NaH_2_PO_4_, 5.8 KCl, 5 EGTA, 0.9 MgCl_2_, 5.6 glucose, and 10 H-HEPES, pH 7.4. After bundle destruction, fresh external solution was given to re-establish the corresponding extracellular ionic environment. Two external solutions were used: 3 mM Ca^2+^ solution containing (in mM) 144 NaCl, 0.7 NaH_2_PO_4_, 5.8 KCl, 3 CaCl_2_, 0.9 MgCl_2_, 5.6 glucose, and 10 H-HEPES, pH 7.4; and 35 mM Ca^2+^ solution containing (in mM) 80 NaCl, 0.7 NaH_2_PO_4_, 5.8 KCl, 35 CaCl_2_, 0.9 MgCl_2_, 5.6 glucose, and 10 H-HEPES, pH 7.4. Only traces with obvious single channel events were included for analyzing.

The sampling rate was 1 kHz for leak current recording, 50 kHz for the IV protocol and current-clamp recording, and 100 kHz for unitary channel recording. The voltage-clamp used a –70 mV holding potential, and the current-clamp was held at 0 pA. Only recordings with a current baseline <20 pA in NMDG solution were used for statistical analysis.

### Hair cell stimulation

The hair bundle was deflected by two types of mechanical stimulus, fluid jet and glass probe. The fluid jet stimulation was as described previously (Beurg et al., 2014). A 40-Hz sinusoidal wave stimulus was delivered by a 27-mm-diameter piezoelectric disc driven by a home-made piezo amplifier pipette with a tip diameter of 3–5 µm positioned 5–10 µm from the hair bundle to evoke maximum MET currents. For glass probe stimulation, hair bundles were deflected with a glass pipette mounted on a P-885 piezoelectric stack actuator (Physik Instrumente, Karlsruhe, Germany). The actuator was driven with voltage steps that were low-pass filtered at 10 KHz. To avoid bundle damage caused by over-stimulation, the glass probe was shaped to have a slightly smaller diameter than the hair bundles, and the stimulation distance was 800 nm for macroscopic current and 100 nm for unitary channel recording.

### Inhibitors, ion substitution, permeability, and perfusion

In Figure 5, DHS, dTC, amiloride, GdCl_3_, and LaCl_3_ were added as calculated to the recording solution (in mM) 144 NaCl, 0.7 NaH_2_PO_4_, 5.8 KCl, 1.3 CaCl_2_, 0.9 MgCl_2_, 5.6 glucose, and 10 H-HEPES (pH 7.4). Dose-inhibition curves were fitted with a Hill equation:

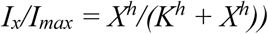

Where *K* is the half-inhibition dose (IC50) and *h* is the Hill slope. *I*_*max*_ is the maximal current in control condition.

In Figure 6, all the ion substitution solutions were derived from a simplified external solution (in mM): 147 NaCl, 1.3 CaCl_2_, 5.6 glucose, and 10 H-HEPES (pH 7.4). In Figure 6A, LiCl and CsCl were 150 mM, completely substituting for NaCl. In Figure 6B, the Ba^2+^ solution was (in mM) 10 BaCl_2_, 137 NaCl, 1.3 CaCl_2_, 5.6 glucose, and 10 H-HEPES (pH 7.4); the Zn^2+^ solution was 75 ZnCl_2_, 75 NaCl, 1.3 CaCl_2_, 5.6 glucose, and 10 H-HEPES (pH 7.4); the Co^2+^ solution was 75CoCl_2_, 75 NaCl, 1.3 CaCl_2_, 5.6 glucose, and 10 H-HEPES (pH 7.4); the Mg^2+^ solution was 150 MgCl_2_, 5.6 glucose, and 10 H-HEPES (pH 7.4); and the Ca^2+^ solution was 75 CaCl_2_, 75 NaCl, 5.6 glucose, and 10 H-HEPES (pH 7.4).

Ca^2+^ permeability was measured by performing whole-cell voltage-clamp recording on P6 OHCs, with intracellular solution contains (in mM): 140 CsCl, 1 MgCl_2_, 0.1 EGTA, 2 Mg-ATP, 0.3 Na-GTP, and 10 H-HEPES, pH 7.2. A voltage ramp stimulation from −120 to 80 mV lasting for 2 seconds was applied to calculate the reversal potential. For measuring Na^+^ permeability, OHCs were perfused with the external solution containing (in mM): 150 NaCl,1.3 CaCl_2_, 5.6 glucose and 10 H-HEPES. For measurement of Ca^2+^ or Mg^2+^ permeability, 150 NaCl with be substituted to 75 Ca^2+^ or 75 Mg^2+^ supplemented with 75 NMDG^+^. In order to eliminate the influence of technical leak, an identical voltage ramp stimulation was applied on each recorded OHC in 150 NMDG. The part of inward current trace was fitted linearly to calculate the voltage value cross point between interest of ion and NMDG solution, which represented the reverse potential of the leak between this ion and Cs^+^. The relative permeability of monovalent cation was calculated as described (Hille)

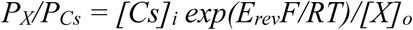

And for divalent cations, the equation was:

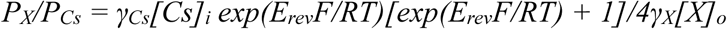

For which γ_Cs_ = 0.70 (Hille), γ_Ca_ = 0.4657, γ_Mg_ = 0.5271 (Rodil and Vera, 2001). *E*_*rev*_ means reversal potential, *F* and *R* mean Faraday constant and gas constant, *T* means absolute temperature. For calculation, 25°C was used as value of room temperature.

For the Ca-NMDG solution in Figure 6E-F, 1 CaCl_2_ was exchanged for 2 NMDG-Cl. For the Na-Ca solution in Figures 6G, 2 NaCl was exchanged for 1 CaCl_2_. The osmotic pressure of each solution was readjusted to 300–320 mOsm/kg with sucrose, and the pH was adjusted to 7.4.

The gravity perfusion system (ALA-VM8, ALA Scientific Instruments, Farmingdale, NY) is controlled manually to switch and deliver solutions. The perfusion tubing and tip were modified as previously reported (Wu et al., 2005). For cochlear tissue, the perfusion tip was placed 2-3 mm from the patched hair cell and the perfusion rate was ∼0.5 ml/min. Extra solution in the recording dish was removed by a peristaltic pump (PeriStar, World Precision Instruments, Sarasota, FL) to maintain a steady liquid level.

### Data analysis

Every experiments contained at least 3 biological replicates and over 10 cell numbers, which were collected at least every 2 weeks within 3 months to keep the stability of a set of data. For certain experiment such as single channel recording, the traces number were over 100. All cell numbers were noted in the figure legends. Multiple recordings from one cell with the identical stimulus protocol were considered as technical replications, which were averaged to generate a single biological replication representing value/data from one cell. Data were managed and analyzed with Excel (Microsoft), Prism 6 (GraphPad Software, San Diego, CA), and Igor pro 6 (WaveMetrics, Lake Oswego, OR). All data are shown as mean ± SEM. Unpaired Student’s t-test was applied to determine statistical significance with two-tailed P values (*p <0.05, **p <0.01, ***p <0.001). Values and N numbers are defined in the figures and figure legends.

## ACKNOWLEDGMENTS

We thank Drs Ulrich Mueller, Anthony Ricci, Bailong Xiao, Xin Liang, Wei Zhang, and members of Xiong laboratory for helpful discussions and critical proof-reading of this manuscript. This work was supported by the National Natural Science Foundation of China (31522025, 31571080, 81873703, and 3181101148), Beijing Municipal Science & Technology Commission (Z181100001518001), and a startup fund from the Tsinghua University-Peking University Joint Center for Life Sciences W.X. is an awardee of the Young Thousand Talent Program of China.

## AUTHOR CONTRIBUTIONS

S.L. did the hair-cell electrophysiology and analyzed data; S-F.W. made the constructs; L-Z.Z. performed the cochlear culture and injectoporation; J.L. performed the hair-cell electrophysiology; C.S., J.C., Q.H., and L.L. conducted the cell culture and molecular experiments; W.X. supervised the project, designed experiments, generated figures, and wrote the manuscript.

---

**Figure S1.**
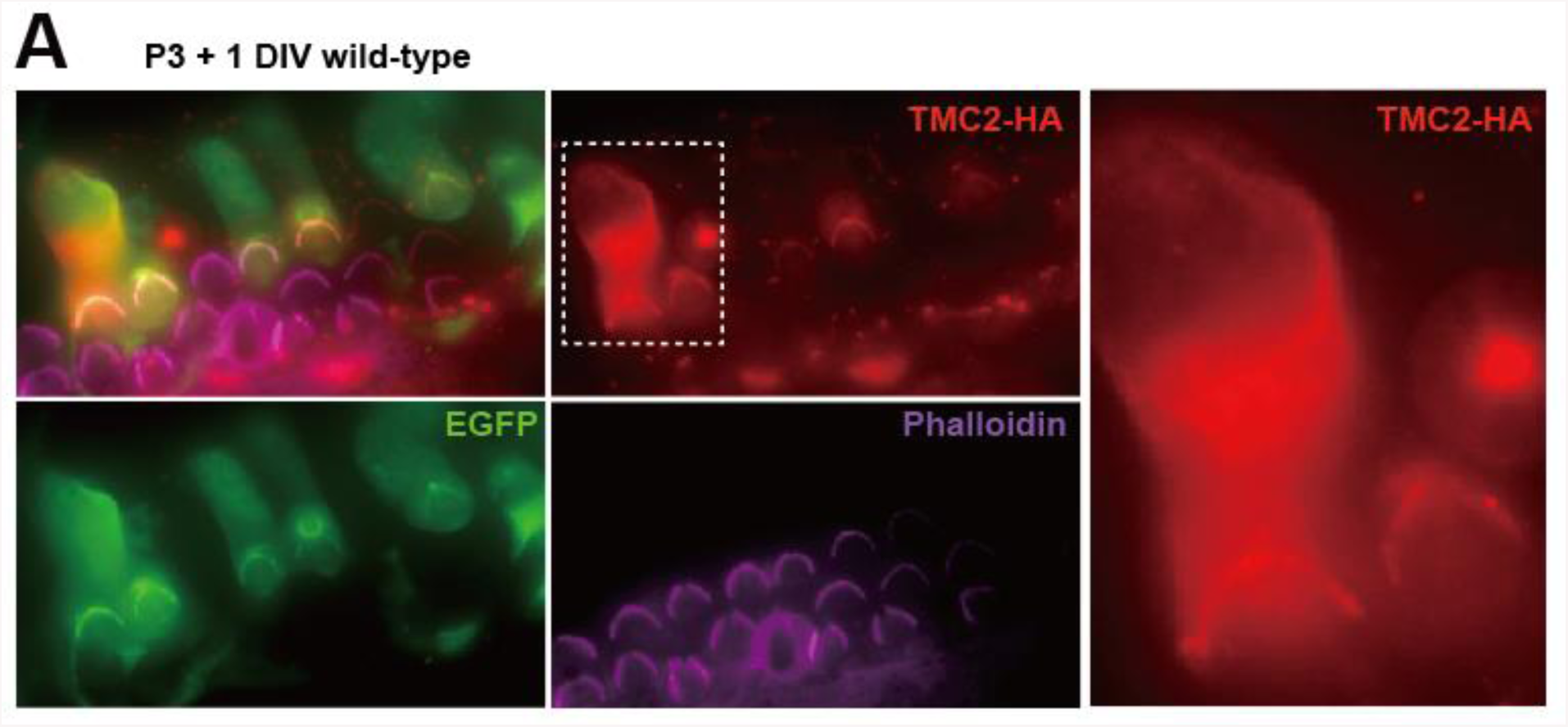
Localization of ectopically expressed TMC2 in OHCs, related to Figure 2. (A) Exogenous expression of TMC2 in P3 + 1DIV OHCs. The OHCs were stained by antibodies to show spatial pattern of TMC2 (by HA antibody, red) and EGFP (by antibody, green). The hair bundle was stained by Phalloidin (magenta). Right: two transfected OHCs were shown enlarged from the white dashed frame.

**Figure S2.**
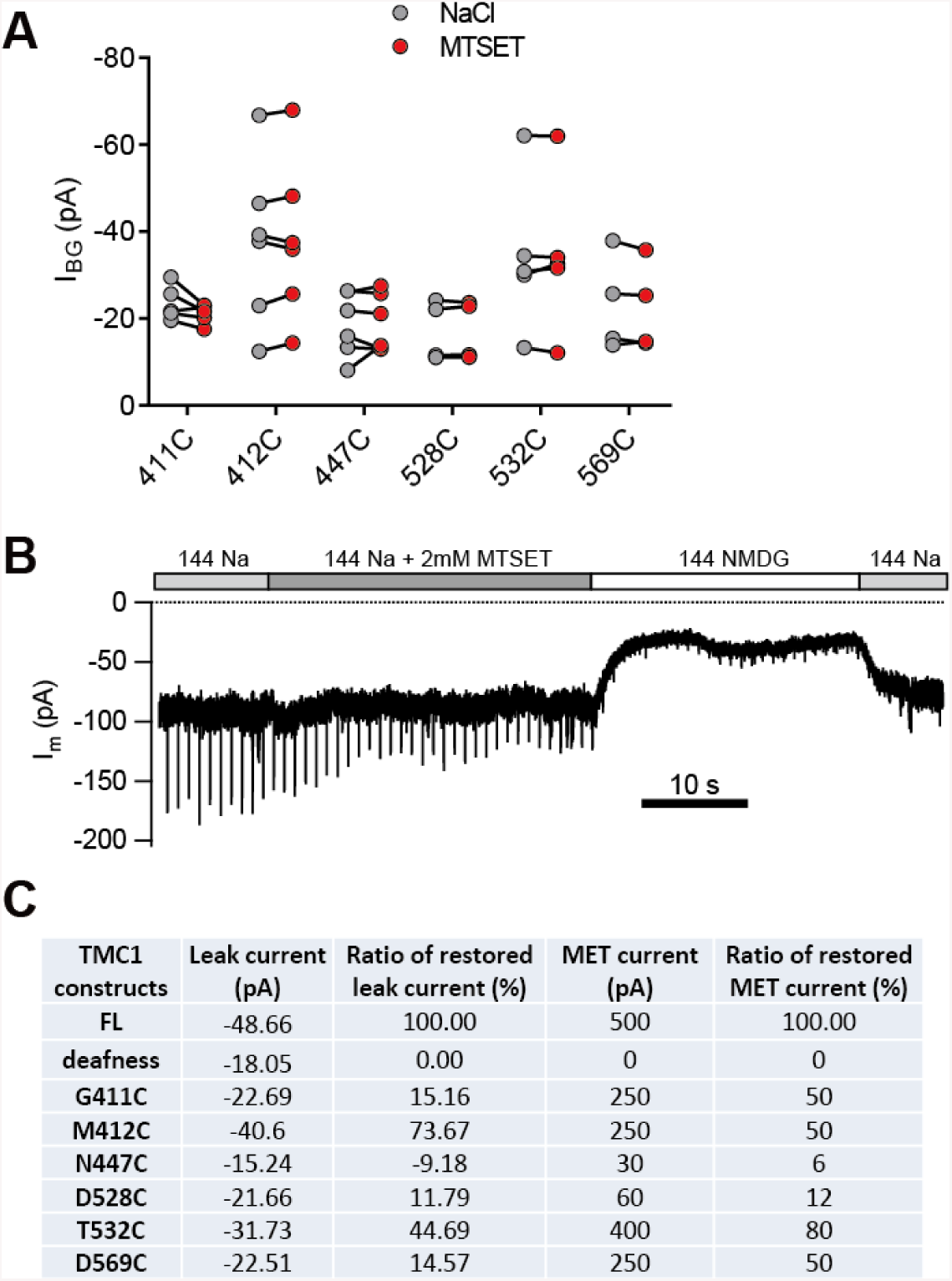
Cysteine substitution in TMC1 affects the MET current and the leak current, related to Figure 4. (A) Plots of amplitude of the background current recorded from *Tmc1*-knockout OHCs expressing engineered TMCs as indicated, before and after MESET treatment. (B) Representative trace of I_m_ recording in a *Tmc1;Tmc2* double-knockout OHC expressing TMC1-M412C. A train of 800 nm step deflection was applied to the hair bundle by a glass probe. (B) Summary of absolute values and normalized ratios of I_Leak_ and I_MET_. The restored MET values of all TMC1 constructs were measured from Pan et al., 2018, excepting that for dn, which was from Kawashima et al., 2011.

**Table S1.**
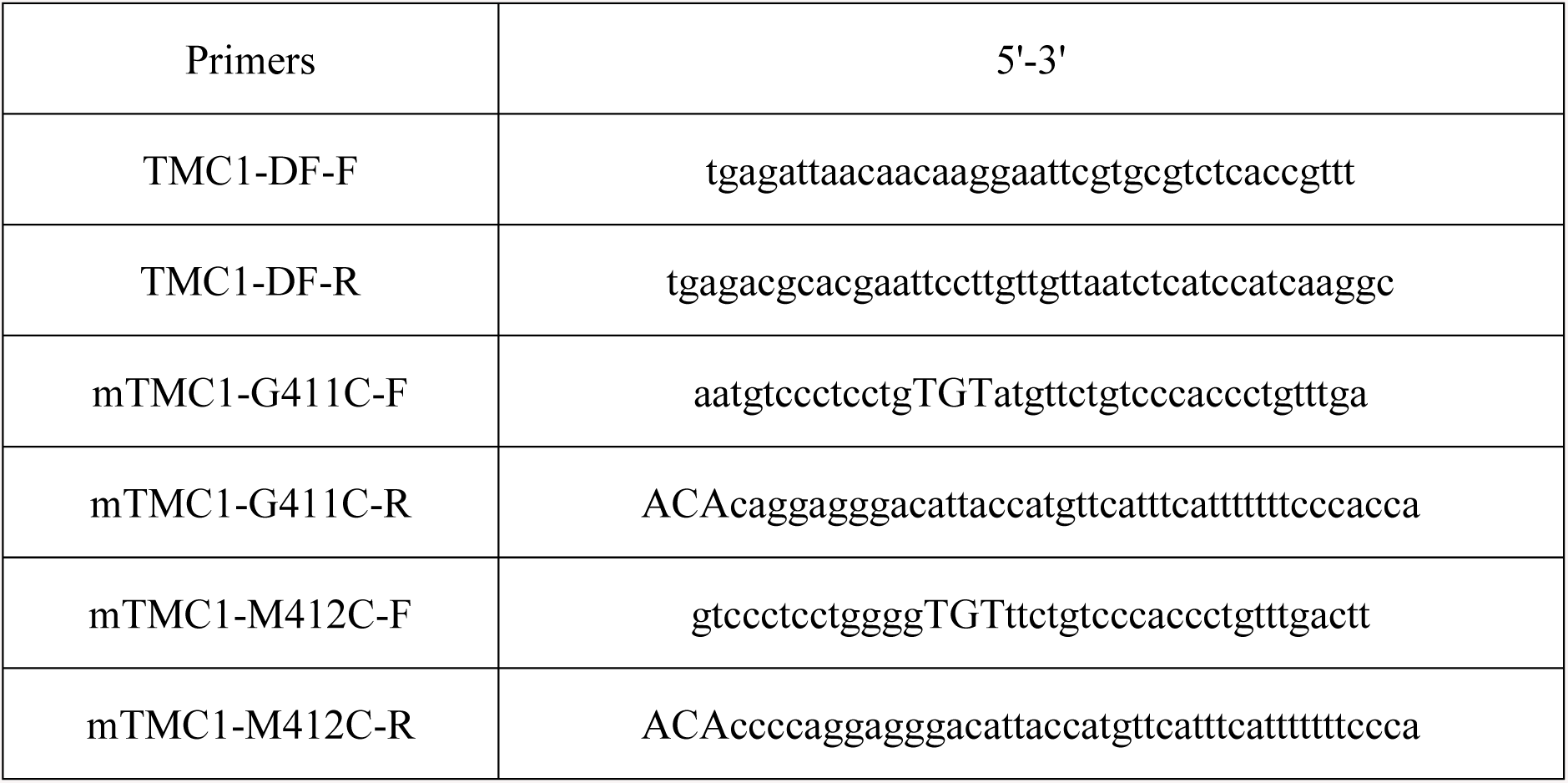
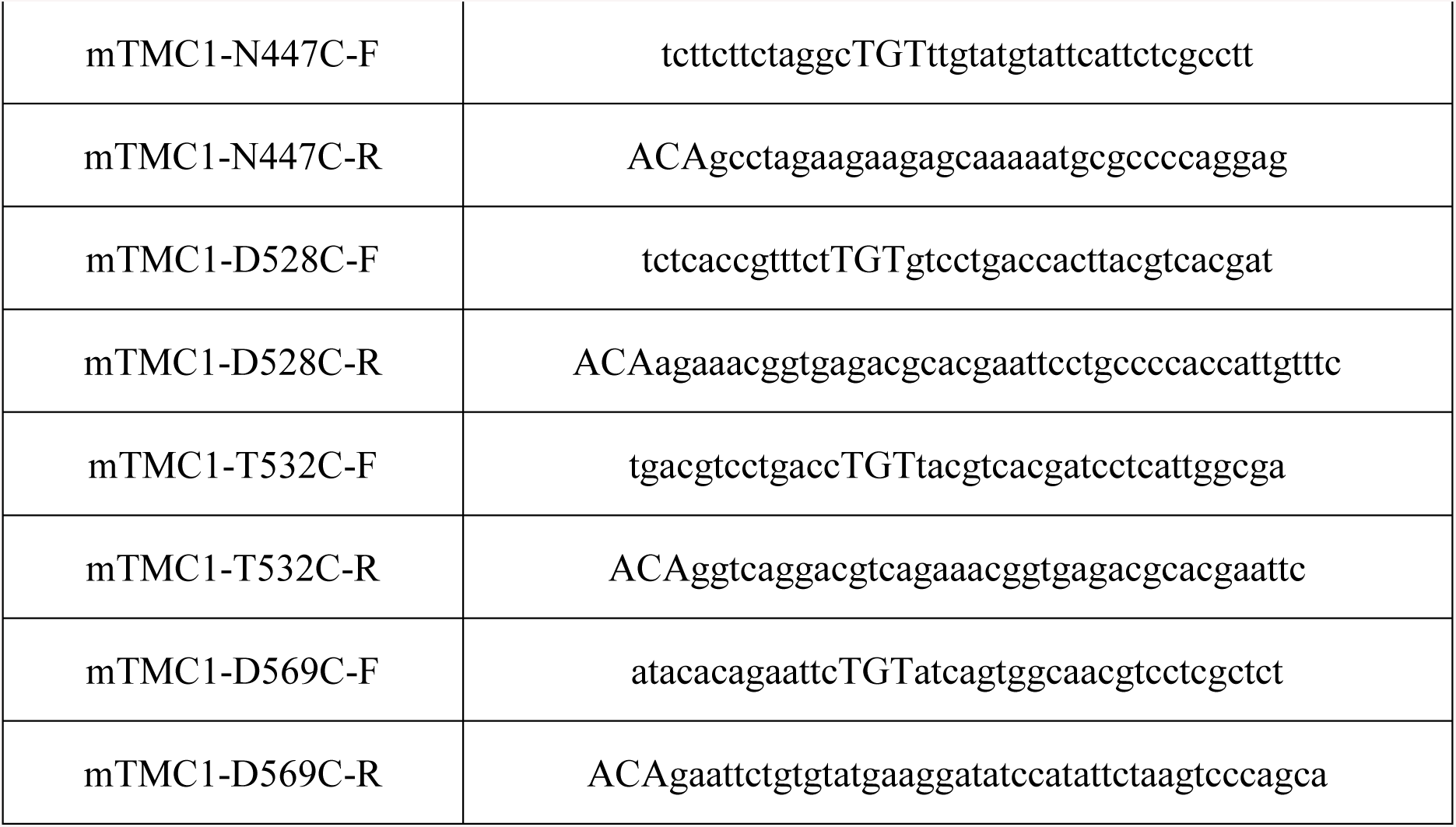
Primers used for generating desired truncation and mutations in mouse *Tmc1* cDNA.

